# Towards mechanistic models of mutational effects: Deep Learning on Alzheimer’s Aβ peptide

**DOI:** 10.1101/2021.12.19.473403

**Authors:** Bo Wang, Eric R. Gamazon

## Abstract

Alzheimer’s Disease (AD) is a debilitating form of dementia with a high prevalence in the global population and a large burden on the community and health care systems. AD’s complex pathobiology consists of extracellular β-amyloid deposition and intracellular hyperphosphorylated tau. Comprehensive mutational analyses can generate a wealth of knowledge about protein properties and enable crucial insights into molecular mechanisms of disease. Deep Mutational Scanning (DMS) has enabled multiplexed measurement of mutational effects on protein properties, including kinematics and self-organization, with unprecedented resolution. However, potential bottlenecks of DMS characterization include experimental design, data quality, and the depth of mutational coverage. Here, we apply Deep Learning to comprehensively model the mutational effect of the AD-associated peptide Aβ_42_ on aggregation-related biochemical traits from DMS measurements. Among tested neural network architectures, Convolutional Neural Networks (ConvNets) and Recurrent Neural Networks (RNN) are found to be the most cost-effective models with robust high performance even under insufficiently-sampled DMS studies. While sequence features are essential for satisfactory prediction from neural networks, geometric-structural features further enhance the prediction performance. Notably, we demonstrate how mechanistic insights into phenotype may be extracted from the neural networks themselves suitably designed. This methodological benefit is particularly relevant for biochemical systems displaying a strong coupling between structure and phenotype such as the conformation of Aβ_42_ aggregate and nucleation, as shown here using a Graph Convolutional Neural Network (GCN) developed from the protein atomic structure input. In addition to accurate imputation of missing values (which here ranged up to 55% of all phenotype values at key residues), the mutationally-defined nucleation phenotype generated from a GCN shows improved resolution for identifying known disease-causing mutations relative to the original DMS phenotype. Our study suggests that neural network derived sequence-phenotype mapping can be exploited not only to provide direct support for protein engineering or genome editing but also to facilitate therapeutic design with the gained perspectives from biological modeling.

## Introduction

As the most common cause of dementia, Alzheimer’s Disease (AD) has been a major research focus in the genomic era, yet effective cures are still lacking^1^. Recent large-scale efforts^2,3^ have sought to characterize the genetic architecture of AD, with a special focus on the discovery of pathogenic variants and disease-relevant genes to elucidate disease mechanisms and, more recently, on leveraging polygenic risk factors for prediction of the disease process at an early point. From a molecular perspective, AD is a neurodegenerative disease characterized by amyloid fiber deposition. Molecular modeling therefore has been performed to identify the critical sequence motifs for peptide aggregation^4^ to guide the discovery of natural or synthetic inhibitors^5^. Despite the achievements, the detailed pathological pathways and mechanisms of AD remain to be elucidated. For instance, the genetic basis of AD is far from being fully understood. Although Genome-Wide Association Studies (GWAS) are reporting associations of an increasing number of genetic variants with the polygenic disease^6,7^, there has been limited understanding of underlying biological mechanisms. In addition, growing evidence indicates that AD may have shared etiologies with other neurodegenerative disorders such as Parkinson’s Disease, raising the question of the trait specificity of relevant molecular mechanisms. Since, in real physicochemical environments, amyloidogenic and other peptides that contribute to the development of AD show a large variational space, exhaustively investigating all possible mutations while capturing peptide phase transition towards mature amyloid deposition is impractical from an experimental or theoretical perspective, especially at atomic resolution^8,9^.

On the other hand, models based on sequence homology detection are often limited by pipelines that require proper substitution matrices, optimized sequence alignment algorithms^10^, and effective scoring functions^11^. Deep Mutational Scanning (DMS) – a high-throughput methodology – has been developed to uncover the functional or biochemical consequences of each possible change at every position in the amino acid sequence^12^. Through large-scale mutagenesis, the effect of mutations (and combinations thereof) may be evaluated in an unbiased manner and disease-causing variants easily identified. However, analysis and interpretation from raw DMS measurements are often biased by many factors, including selective fragments (instead of complete sequence), limited combinations of single mutations (from a combinatorially astronomically large set of complex mutations), and experimental errors (e.g., batch effects or background noise). Deep learning presents new opportunities for generating a high-resolution atlas of the sequence-phenotype relationship, i.e., establishing neural network (NN) derived models of mutational effects on biochemical phenotypes.^10,13^. Utilizing a previously reported supervised learning framework^14^, we developed NN-derived models of SARS-CoV-2-related biochemical phenotypes, namely, binding affinity and protein expression. Critical mutations in the virus and the host were identified via our NN-derived, mutationally-determined phenotypes, and their biological consequences were further determined. The findings were confirmed in independent experimental data^15^. Notably, Convolutional Neural Networks (ConvNets), showed much stronger prediction performance than linear regression or single-layer Fully-Connected Neural Networks (FCN).

Among all Aβ isoforms, Aβ_42_ is the most aggregation-prone, predisposing to Alzheimer’s disease via self-recognition and replication^16^. DMS experiments have been performed on Aβ_42_, aiming to elucidate the genetic landscape as well as the molecular determinants of aggregation, indexed by either nucleation^3^ or solubility^17^ for each residue. Here, we considered seven neural network architectures to investigate the sequence-phenotype map, including sequence Convolutional Neural Networks (seq-CNNs), Recurrent Neural Networks (RNNs), and Graph Convolutional Neural Networks (GCNs). Importantly, we designed neural networks to enable extraction of biological insights during the model training. Beyond functioning as vectorized input features^18^, protein structures could be integrated into the model architecture, thus not only facilitating the prediction of biochemical phenotypes but also modeling biological phenomena towards mechanistic interpretation.

## Results

### Deep learning framework and performance on aggregation-related biochemical phenotypes

Seven network architectures implemented here fall into three classes: 1) Sequence-based: This class includes two seq-CNNs, i.e., CNN-1D and CNN-2D, and an RNN architecture with Long Short-Term Memory (LSTM) cells. 2) Sequence- and structure-based: This class includes two GCNs, i.e., Single Weight (GCN-SW) and Average Node (GCN-AVE). 3) Baseline: This class contains Linear Regression (LR) and Multilayer Perceptron (Fully-Connected Neural Networks [FCN]). Additional details on the network architectures are found in **Methods**. Briefly, 20 amino acids and 1 stop codon comprised the essential “vocabularies” in the RNN model, wherein sequence-phenotype profiles were tokenized and padded. In the case of CNNs, GCN and FCNs, besides the Identity Descriptor (one-hot encoding), we further utilized the Amino Acid Based Descriptor, AAindex^19^, which had been generated from the projection of 566 intrinsic amino acid physicochemical properties (e.g., hydrophobicity, long-range non-bonded energy per atom, solvent-accessible surface area) to a 19-dimensional vector space^19^ (Figure 1A). Compared to standard CNNs, GCNs apply convolution operators to Generative Graph Architectures (GGAs) generated from protein structural coordinates, such that prediction occurs at the node/residue level^20,21^ (Figure 1A inset). Three aggregation-related biochemical phenotype measurements were adapted from two separate DMS studies, i.e., nucleation^3^, solubility^17^, and ‘synonymous’^17^, directly or indirectly reflecting Aβ_42_ aggregation.

**Figure 1.**
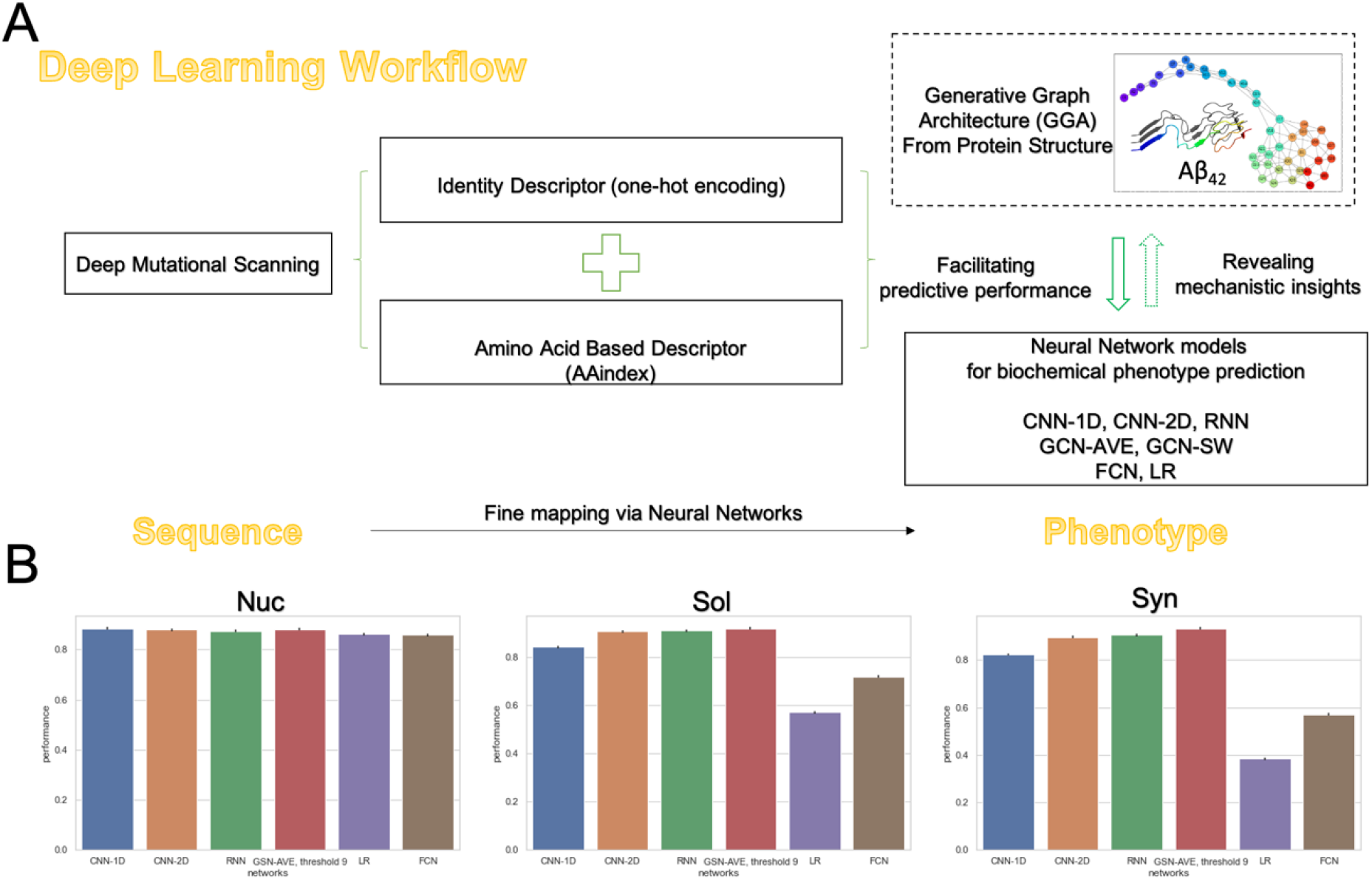
Neural networks for mapping the sequence – biochemical phenotype relationship for Aβ_42_. **A**. Our Deep Learning pipeline is built on the sequence-phenotype input from Deep Mutational Scanning (DMS). The pipeline leverages the Identity Descriptor (one-hot encoding) and the Amino Acid Based Descriptor (representing 566 intrinsic physicochemical properties of each amino acid). Generative Graph Architectures (GGAs) generated from protein structural information (i.e., atomic coordinates) are integrated into select neural network models to facilitate phenotype prediction. In addition, the neural network architectures along with the derived phenotype from each of the predictive models were used to extract mechanistic insights underlying established biological understanding. **B**. For each model trained in a training set, Spearman coefficient correlation between the observed and predicted phenotype in an independent test set was used to evaluate the prediction performance (**Methods**). The three phenotypes modeled here were nucleation (Nuc), solubility (Sol), and ‘synonymous’ (Syn). Model performances were compared across six representative neural network models on each phenotype. ConvNets and RNN demonstrated robustly strong performance for all phenotypes, with Graph Convolutional Neural Networks outperforming all models, while the baseline models (LR and FCN) performed well only for the phenotype with the largest (two orders of magnitude greater) training data.

We hypothesized that a well-modeled fibril conformation might provide crucial insights into aggregation, enhancing downstream mechanistic analysis. The prediction performance of each neural network model developed in a training set was evaluated in an independent test set (**Methods**) using the Spearman coefficient correlation between the predicted and observed phenotype (Figure 1B, Table S1). For each model, the standard error was estimated from the application of bootstrap (*n* = 100). GCNs consistently showed superior performance (correlation ∼ 0.9) for all phenotypes despite varying the input GGAs, which were derived from three different Aβ_42_ conformations (**Methods**). RNN-LSTM also achieved high performance, illustrating its distinctive advantages processing the sequential input. In contrast, LR and FCN achieved good prediction for the nucleation phenotype but showed degraded performance for the other two phenotypes. Given the nearly 14,000 samples collected for the “nucleation” DMS experiments, two orders of magnitude larger than the sample sizes for solubility and “synonymous” (because of missing data for both phenotypes), the performance drop for the baseline architectures could be the result of differential power or the quality of the training data. The decline may also be due to the presence of two distinct aggregation pathways being implicated by the DMS experiments, as previously proposed^3^, which we will touch on later.

### Extracting mechanistic insights from sequence-based neural networks to model biochemical phenotype

We explored the architectures from sequence-based models, wherein information is derived strictly from the sequence-phenotype relationship, to gain insights into the importance of local sequence information at each residue for the phenotype. To investigate the hyperparameters as well as interpret the information learned from neural networks, we implemented a simple architecture comprising only one dilated CNN-1D layer (**Methods**) to predict the solubility phenotype. Specifically, a dilated kernel of size *k*′ would scan the residues using intervals – adjusted by a dilation rate *α* ∈ ℕ, *α* ≥1 – as opposed to local motifs, to cover a possibly larger sliding window containing *non-adjacent* neighboring residues (Figure 2A), whose effect on performance can be quantified. For *α* > 1, the impact of the residues skipped during the dilated convolution on the prediction performance can be evaluated through comparison with the performance in the absence of any dilation (i.e., *α* = 1). The modified kernel size *k*′ was defined as follows: *k*′ = *α* ∗ (*k* − 1) + 1, from the size *k* of the actual kernel altered by a range of values for the dilation rate *α* (Figure 2B, Table S2). In the absence of dilation, the model architecture reduces to the conventional ConvNets, wherein the optimized kernel size is obtained through hyperparameter tuning. Different kernel sizes were determined while varying *α* and fixing *k*′. For each *k*′, the model performance showed a significant boost, doubling or tripling the actual kernel size *k*. In contrast, for each *k*, e.g., 3, 5, although a larger dilation rate *α* yielded a larger *k*′, i.e., a larger convolutional window, the prediction performance was not necessarily improved and was often degraded (Table S2). These observations strongly imply that local information and neighboring interactions (for example, involving the residues skipped during the dilated convolution) have a substantial impact on the prediction performance. The use of a larger dilation rate, despite the larger convolutional window, neglects such local information and neighboring interactions that may be critical to the predicted solubility. Indeed, from a biological perspective, a few core fragments (corresponding to the absence of dilation, *α* = 1) of Aβ_42_ mediating the overall hydrophobicity of the peptide have drawn significant attention from molecular studies^4,8^, as these fragments are closely related to inter-peptide interactions.

**Figure 2.**
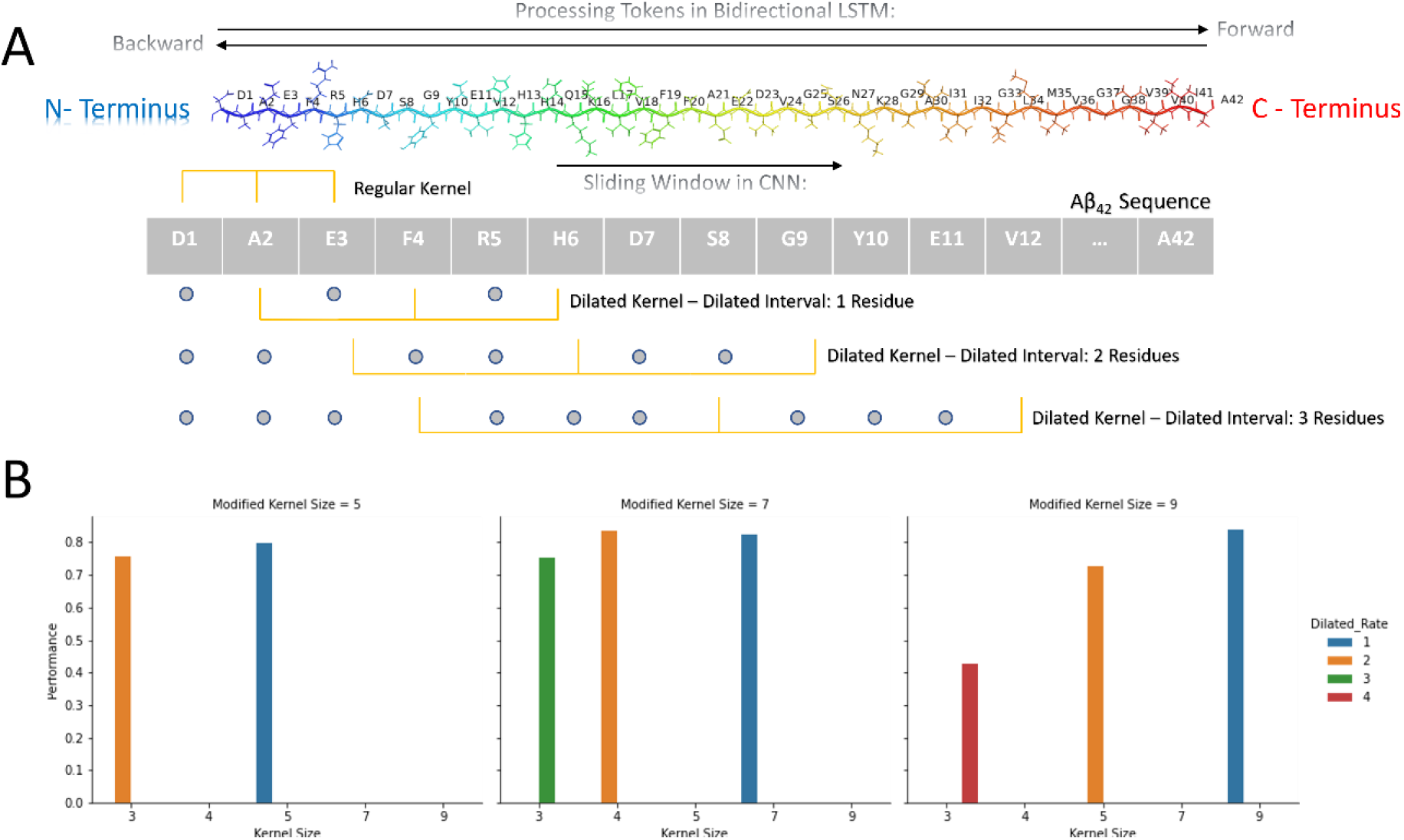
Architectures and performance of sequence-based models. **A**. Aβ_42_ is colored with the “rainbow” scheme from the N-Terminus (blue) to the C-Terminus (red) end. Each amino acid is represented as a one-letter code. In a Recurrent Neural Network (RNN) LSTM architecture, connections between nodes constitute a directed graph along a sequence of time steps, with the LSTM providing a solution to the vanishing gradient problem associated with a RNN (**Methods**). While bidirectional LSTM enables the network to exploit context on both sides of each position via two independent RNNs (Forward and Backward), 1D Convolutional Neural Network (CNN-1D) slides from the N- to the C-terminus with regular and dilated kernels. A dilated kernel covers a larger convolutional window with selected non-adjacent neighboring residues, enabling evaluation of their contribution to prediction performance. Here, interval residues are shown as gray dots. **B**. Model performance of one-layer CNN-1D with varying modified kernel size *k*′or dilation rate *α* ∈ ℕ, *α* ≥1 (Table S2). For a model trained in a training set, Spearman coefficient correlation between the observed and predicted phenotype in an independent test set was used to evaluate the prediction performance (**Methods**). Actual kernel size *k* had a substantial impact on prediction performance with (*α* > 1) or without (*α* = 1) dilation. Increasing the dilation rate with the same kernel size yielded a larger convolutional window, but performance was often degraded (Table S2), indicating the importance of local information and neighboring interactions.

We implemented the CNN-1D architecture by adapting ProtCNN^22^, which wraps dilated layers in residue blocks and was originally used for computationally efficient protein classification (**Methods**). Protein sequence inputs could also be trained using a CNN-2D architecture by extending the dimension^14^. Compared to a 1D architecture, higher-dimensional architectures displayed satisfactory performance for all the phenotypes with a simple architecture setup (Figure 1B, Table S1). On the other hand, good performance was also achieved from RNN-LSTM models even with only the Identity Descriptor (Table S1). Bidirectional LSTMs, by default, train two LSTMs, the first on the input sequence and the second on a reversed copy, providing contextual information to the network^23^. While we had highlighted the significance of Amino Acid Based Descriptor in a previous study^15^, here we found that the choice of model algorithm had an even stronger impact than the deployment of features (Table S1) as claimed in a recent study^18^.

### Extracting mechanistic insights from sequence- and structure-based neural networks to model biochemical phenotype

In this section, we present results on the contribution of structural features to the neural network performance. The atomic coordinates of the protein residues were used to generate a Generative Graph Architecture (GGA), wherein nodes represent residues and edges correspond to inter-residue connections (representing, for example, covalent bonds, hydrogen bonds or van der Waals interactions) under a pre-defined Euclidean distance threshold or physical interaction radius (Figure 3A, **Methods**). By varying the GGA input into a neural network, we could evaluate the contribution of specific residues (nodes) or interactions (edges) to differential prediction performance. In contrast to the Node Average model (GCN-AVE), the Single Weight model (GCN-SW) sums up the contributions of all nodes within a convolutional window without explicitly differentiating the node types and averaging the contributions^20^ (**Methods**). We implemented GCN-SW as a model architecture to test against nucleation, which directly reflects the aggregation tendency of the mutants at each residue position of Aβ_42_. We set the number of layers and filter size to 1, so that the prediction performance of GCN-SW is primarily determined by the GGA inherited from the protein structure input, i.e., without any additional optimization on the network architecture.

**Figure 3.**
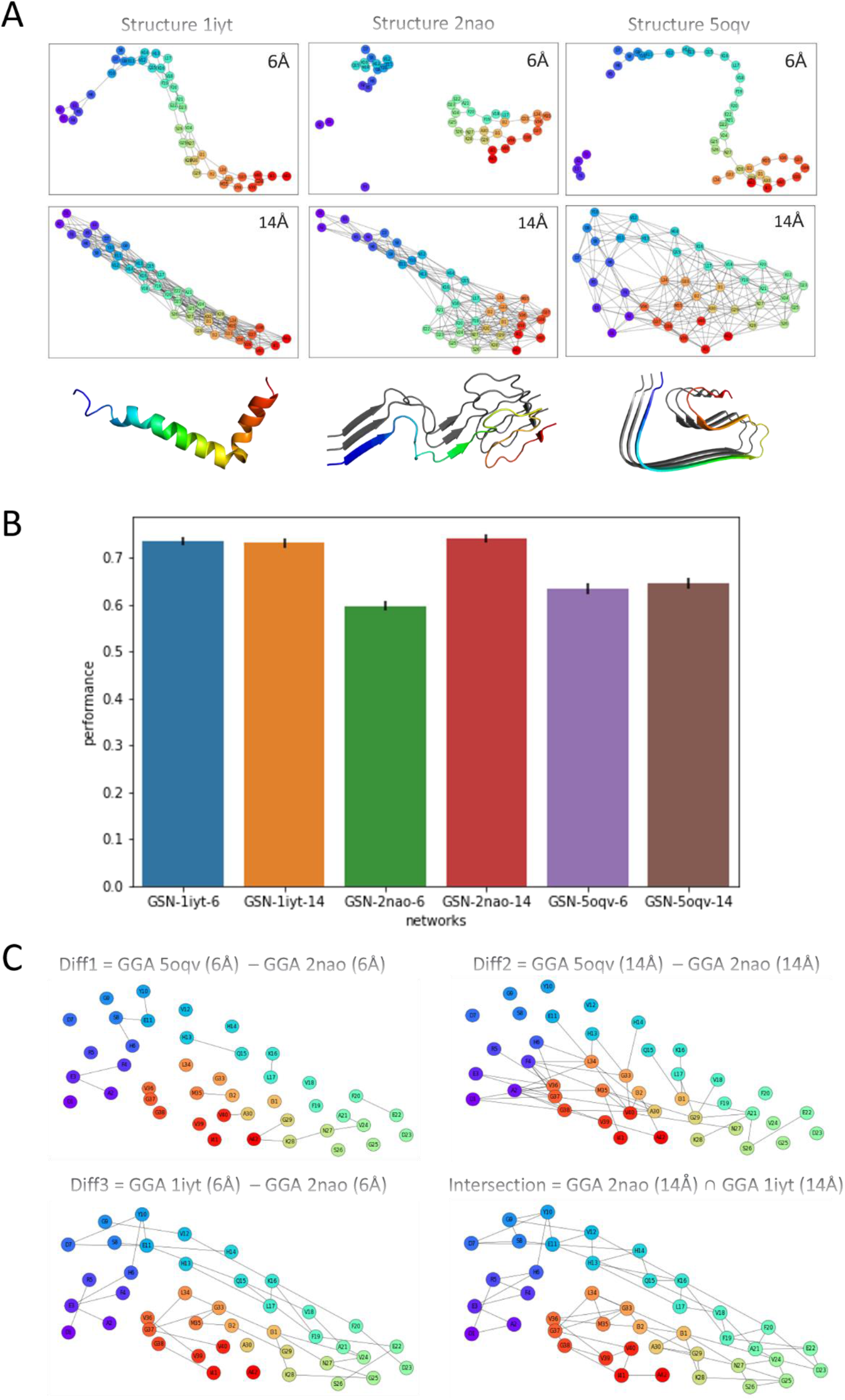
Biological inferences from Single Weight Graph Convolutional Neural Networks (GCN-SW). **A**. Generative Graph Architectures (GGAs) generated from three Protein Data Bank structures (determined by NMR spectroscopy or cryo-electron microscopy), 1iyt, 2nao and 5oqv, with two representative Euclidean distance thresholds, 6Å and 14Å. Each residue is represented by a node while the presence of a connecting edge between nodes is determined by whether the Euclidean distance between the residues (calculated from the atomic coordinates of the corresponding *β*-carbons) is within the chosen distance threshold. Thus, chemically, the edges may represent intermolecular interactions, including covalent bonds, hydrogen bonds or van der Waals interactions between consecutive residues or non-consecutive residues. Nodes from GGAs were mapped to the protein sequence (with each node represented by a one-letter code) and colored with the “rainbow” scheme from the N-terminus (blue) to C-terminus (red), which is here also reflected in the protein tertiary structure (in third row). Note the GGAs for the alpha-helix-rich 1iyt resembled the connected peptide backbone while 2nao and 5oqv, both in mature fibril conformations, displayed multiple connected components at 6Å. Diverse structural inputs of protein atomic coordinates lead to contrasting GGAs. **B**. Model performance of GCN-SW (filter size=1, number of layers=1) in predicting nucleation based on the six GGAs, implemented as described in the text. **C**. New graph networks derived from “Graph Operations” (Difference and Intersection; **Methods**) on the initial six GGAs. The top 3 nodes/residues with high centrality were identified in the “Diff3” and “Intersection” GGAs.

Monomeric Aβ_42_ is intrinsically disordered with varying conformations under changing physicochemical conditions. Three structures (determined by NMR spectroscopy or cryo-electron microscopy) were retrieved from the Protein Data Bank with the following PDB IDs: 1iyt(NMR)^24^, 2nao(NMR)^25^ and 5oqv(cryo-EM)^26^ (Figure 3A for protein snapshots). Specifically, the structure 1iyt had been characterized in monomer form under the aqueous solutions of fluorinated alcohols – highly hydrophobic environments – thus is rich in helices. In contrast, the structures 2nao and 5oqv had been characterized in mature fibril form, yet the conformations substantially differed in the presence of stacked layers of beta sheets. We used the protein structure coordinates to generate GGAs with varying number of edges and node degrees, as functions of the chosen Euclidean distance threshold. Varying this threshold over values ranging from 4Å to 15Å (Figure S1), we chose 6Å and 14Å as representative thresholds to illustrate the impact of altering the input structural features in GCN-SW on model performance (Figure 3B) and to evaluate the contribution of specific residues or interactions.

In the three-dimensional space of protein atomic coordinates, the 6Å threshold captured, with high fidelity to the 1iyt structure, the local connections among the residues, generating a connected graph highly resembling the peptide backbone (Figure 3A). However, as the 1iyt structure is alpha-helix-rich with a compact conformation, quite a few remote connections were also retained (at the same distance threshold) in the resulting GGA, leading to relatively improved prediction performance (Figure 3B). On the other hand, although both the 5oqv and 2nao structures originated from mature fibril conformations, the GGA from 5oqv yielded improved prediction performance relative to 2nao (Figure 3B). A direct comparison between the two GGAs, which we formalized into a “Graph Operation” for its generalizability (**Methods**), revealed that a few edges/connections, including (A2-E3-F4, H6-S8-E11-Y10, H13-Q15, K16-L17, F20-E22, Q29-A42-K28-N27-V24-A21), were captured only in the 5oqv GCA (as shown in “Diff1” in Figure 3C). These edges are critical for the beta sheet alignments (as can be seen in the snapshots of the fibrils), highlighting the strong correlation between the input protein atomic structure for the neural network model (GCN-SW) and the predicted phenotype (nucleation).

On the other hand, as expected, the larger 14Å threshold captured longer-range inter-residue connections, thus forming more complex graph networks. Reasonably good prediction performance was achieved from the structures 1iyt and 2nao but not from the structure 5oqv (Figure 3B). To investigate the poor performance from the latter, we compared the GGAs derived from the mature fibrils (2nao and 5oqv), again via a Graph Operation. It would appear that the connections uniquely present in the structure 5oqv, most of which connected the N- and C-terminus residues (see connections between blue and red nodes in “Diff2” in Figure 3C), were not substantially contributing to the prediction of the nucleation phenotype. From a structural perspective, interactions among the N- and C-terminus residues would disfavor the phase transition towards the formation of Aβ_42_ fibrils. Notably, for increasing values of the distance threshold, the resulting GGAs of the structure 2nao showed substantially boosted prediction performance (Figure 3B).

Here, we further illustrate the important structural insights that can be universally extracted from a Deep Learning analysis on GGAs through the aforementioned Graph Operations. A Graph Operation-based perturbative approach (**Methods**) developed here has the benefit of identifying relevant features – considered jointly rather than in isolation – by capturing their effects across empirically-observed structures. Let us consider the ‘difference’ in GGAs between the structures 2nao and 1iyt at the 6Å threshold and also the ‘intersection’ of the GGAs of the two structures at the 14Å threshold. The resulting networks illustrate the connections among the C-terminus residues. The nodes of high centrality (top 3) were identified within the two derived GGAs, namely, the “Diff3” GGA (identifying the residues E11, K16, I31, E3, Y10, Q15, F19, V24, N27, G37) and the “Intersection” GGA (identifying the residues F20, V24, I31, N27, Y10, E11, H13, K16, A30, I32, V36, V39) (see Figure 3C for “Diff3” and “Intersection”). Besides these high-centrality nodes/residues in the C-terminus contributing to the beta sheet formation, several high-centrality nodes/residues in the N-terminus, including E11, K16, Y10, and V24, are also of functional interest. Both the neural network prediction and the biological context imply that, for each such residue: 1) the magnitude of the nucleation effect of a mutation at the node/residue is relatively strong, e.g., E11 (Figure 4A & 4B); or 2) the residue has an edge/interaction with other residues, which contributes to differential predictive performance on nucleation (Figure 3B). Our results show not only that modeling via GCNs facilitates insights into mutational effects on the phenotype, but also that the network architecture itself could be designed to infer biological insights (in this case, on the centrality of specific residues [nodes] and the role of specific interactions [edges] for nucleation).

**Figure 4.**
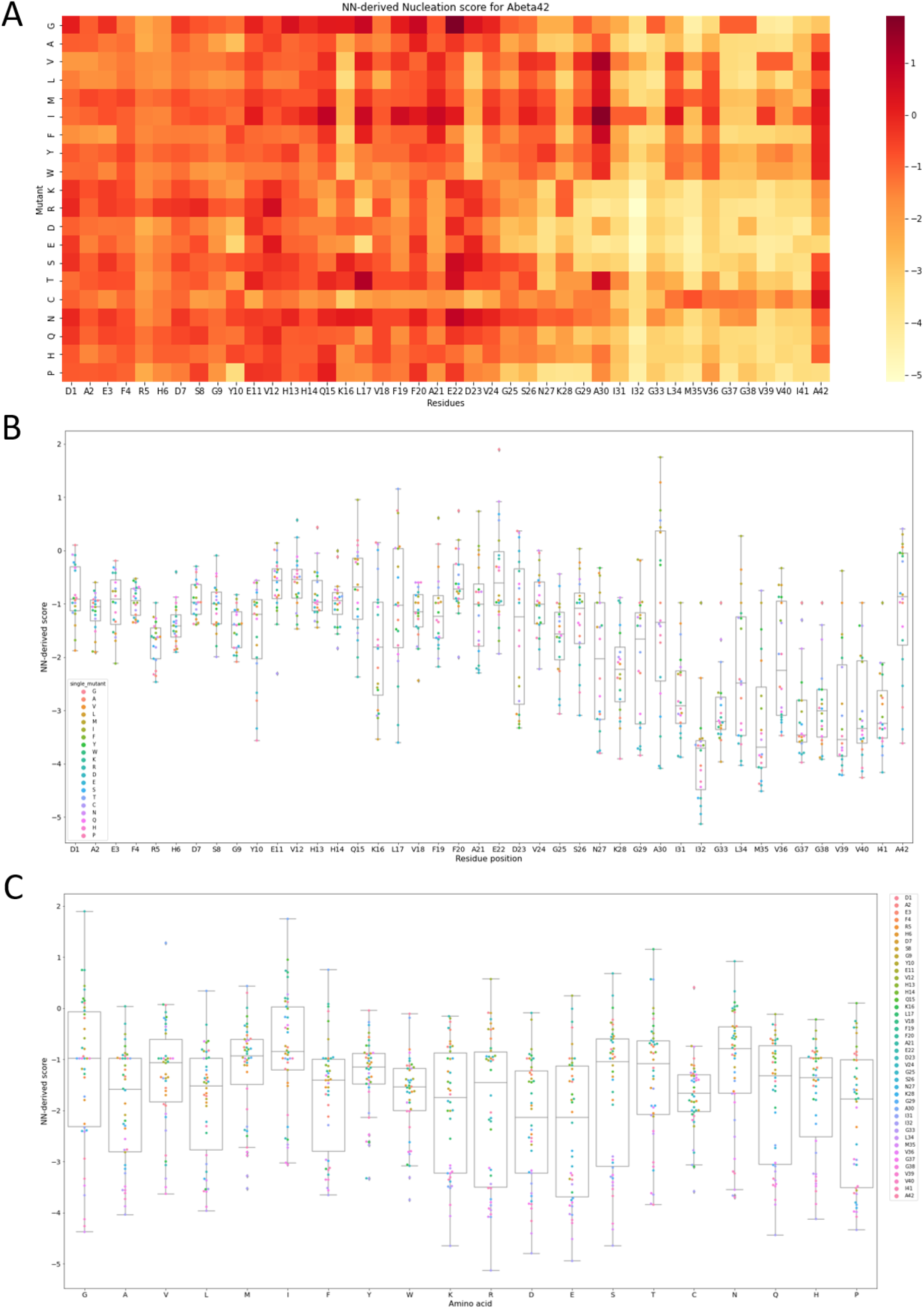
Mutational effects on nucleation in Aβ_42_. **A**. Heatmap and **B**. boxplots generated from our NN-derived nucleation score show the effects of all possible mutations at every single position along the sequence. One benefit of the NN-derived map is the imputation of missing values, which ranged up to 55% at “gatekeeper” residues (as defined by Seuma *et al*.). The C-terminus (S26-A42) had significantly lower NN-derived nucleation than the N-terminus (Mann-Whitney U test, *p* < 2.2×10^−16^). The threshold for the NN-derived score that discriminates positive or negative nucleation effect, equal to the mode of the distribution, is located at −1.041. **C**. Mutational effects for each of 20 amino acids (x-axis) are summarized for all positions in the Aβ_42_ peptide using the NN-derived, mutationally-determined nucleation score (y-axis).

### Mutationally-determined nucleation and disease consequences of mutations

Three causative genes^27^ – Amyloid Protein Precursor (*APP*), Presenelin-1 (*PSEN1*), and Presenelin-2 (*PSEN2*) – and the genetic risk factor *APOE*ε4^28^ are among the well-known associations with AD. *APP* encodes the Aβ precursor protein, which is processed by the β-secretase and the γ-secretase resulting in the Aβ peptide. The presenelins (*PSEN1* and *PSEN2*) comprise the catalytic subunit of the γ-secretase complex. Missense mutations in *PSEN1* are the most common cause of early-onset familial AD (fAD), resulting in a relative increase in the isoform ratio Aβ_42_/_40_Aβ ^29^. In addition, the deposition of Aβ_42_ may be an early preclinical event in *PSEN1* mutation carriers^30^.

We implemented neural networks to generate mutationally-determined nucleation phenotypes and impute a substantial proportion of missing values at key residues (ranging from 30% to 55%), thus completely covering all possible mutations along the Aβ_42_ sequence (Figure 4A), i.e., overcoming the limited and incomplete characterization (due to missing data) from the original DMS experiments by Seuma *et al*.^3^. For ease of presentation, in the results reported below, we used the nucleation phenotype derived from GCN-AVE (with number of layers=2, filter size=32, and threshold=9Å; Table S1), which showed robust prediction performance across all phenotypes.

The finding from the NN-derived nucleation scores on the significant difference (Mann-Whitney U test, *p* < 2.2×10^−16^) between the N-terminus and the C-terminus (S26-A42) (Figure 4A) is consistent with the analysis of the original DMS measurements^3^. This pattern is also reflected in the bimodal empirical distribution of the NN-derived scores (Figure S2). In addition to imputing 40% of the mutagenesis data (comprising the missing data), the NN-derived phenotype, determined or defined as such solely by mutational effects, should be less affected by technical confounders (e.g., batch effects) in downstream analyses. We further illustrate this point here. Mutations in the C-terminus were substantially enriched for decreasing nucleation (Figure 4B), with only 20% in the region increasing nucleation. This observation is not surprising, as C-terminus residues are essential building blocks for beta sheet stacking, so that changing the block’s composition would likely disrupt the alignment^5^. However, A42, which closely associated to the *PSEN1* gene, is a notable exception within the C-terminus, with more than half of the mutations (65%) above our NN-derived discriminating threshold (Figure 4B), which is defined as the mode of the distribution of the NN-derived phenotype (Figure S2A). This outlier pattern for A42 within the C-terminus is clear from our NN-derived nucleation profile (Figure 4A), but not so clear with the original DMS nucleation data, with 55% of the mutation data missing at this position. The previously reported “gatekeeper” residues (at which, by definition, *at least* as many mutations increased as decreased nucleation) from Seuma *et al*., namely D1, E3, D7, E11, L17, E22 and A42, were accurately, and indeed, more definitively, captured by our NN-derived map (Figure 4A & 4B), likely due to the proportion of missing values in the original DMS data at these positions (from 30% to 55%). Moreover, although most mutations were benign or nontoxic, we found that a mutation to Isoleucine (I) or Asparagine (N) could be highly pathogenic at multiple positions (Figure 4C). Taken together, these results show that the NN-derived nucleation was providing significantly greater support than the original DMS phenotype for the critical roles of key residues in Aβ_42_ aggregation.

### Disease mutations and neural network derived phenotype

We investigated 11 disease mutations in Aβ_42_ (https://www.alzforum.org/mutations/app) that are reported to cause dominantly inherited fAD, comparing the effects from the original DMS characterization and from our NN-derived phenotype. For the comparison, single mutations from both datasets were mapped and processed with quantile normalization, thus sharing the same adjusted discriminating threshold (Table 1) equal to the mode of the distribution of the quantile-normalized phenotype (Figure S2B). Agreement with the known disease-causing mutations was found for 10/11 vs 9/11^3^ for our NN-derived vs the original DMS phenotype, respectively. Notably, the NN-derived phenotype yielded the correct prediction for D7H, where the original DMS phenotype failed. Collectively, the results from the NN-derived phenotype showed improved resolution for identifying the known disease-causing mutations.

**Table 1.** Predicted mutational effects for 11 key variants in dominantly inherited familial Alzheimer’s Disease (fAD)

|  | Effect from normalized DMS score | Predicted effect from normalized NN-derived score | NN-derived fAD class | Agreement with actual observations |
| --- | --- | --- | --- | --- |
| A2V | -0.942261566 | -1.11187735 | Recessive | Yes |
| H6R | 0.331734573 | 0.56268832 | Dominant | Yes |
| D7H | 0.61278933 | -0.038973138 | Recessive | Yes |
| D7N | 0.4526346 | 0.598398457 | Dominant | Yes |
| E11K | 0.34264462 | 0.240375209 | Dominant | Yes |
| A21G | 0.35454437 | 0.20493074 | Dominant | No |
| E22G | 2.722336826 | 2.72233683 | Dominant | Yes |
| E22K | 0.740425073 | 0.80116783 | Dominant | Yes |
| E22Q | 2.321731858 | 0.64342195 | Dominant | Yes |
| D23N | 1.268075687 | 1.24206492 | Dominant | Yes |
| A42T | 0.72504662 | 0.758015702 | Dominant | Yes |
| *Threshold that discriminates positive or negative nucleation effect ($\Delta$) has been adjusted and aligned after quantile normalization (Figure S2):<br>Threshold for Raw DMS score: 0 -> -0.5844597<br>Threshold for NN-derived score: -1.041383 -> -0.5844597 | | | | |

### Feature attribution

For improved interpretation of the model on nucleation, we applied the “Integrated Gradients” (IG) method^31^, calculating the normalized feature attribution (Figure 5) for each residue along a mutated sequence. Feature attribution allows systematic examination of the contribution of input features to the prediction. We chose the IG approach because, in contrast to DeepLift^32,33^ and Layer-wise relevance propagation^34^, it combines two (desirable) characteristics, i.e., implementation invariance and sensitivity, of attribution methods. The Aβ_16-22_(KLVFFAE) fragment, which is critical for self-assembly, was found to be highly relevant based on the IG-based feature attribution. In addition, key residues identified through Graph Operations (Figure 3C), including E3, V36, and V39, showed high IG-based feature attribution in the sequence-based CNN-1D model, confirming the inference from Graph Operations on empirically-observed protein structures.

**Figure 5.**
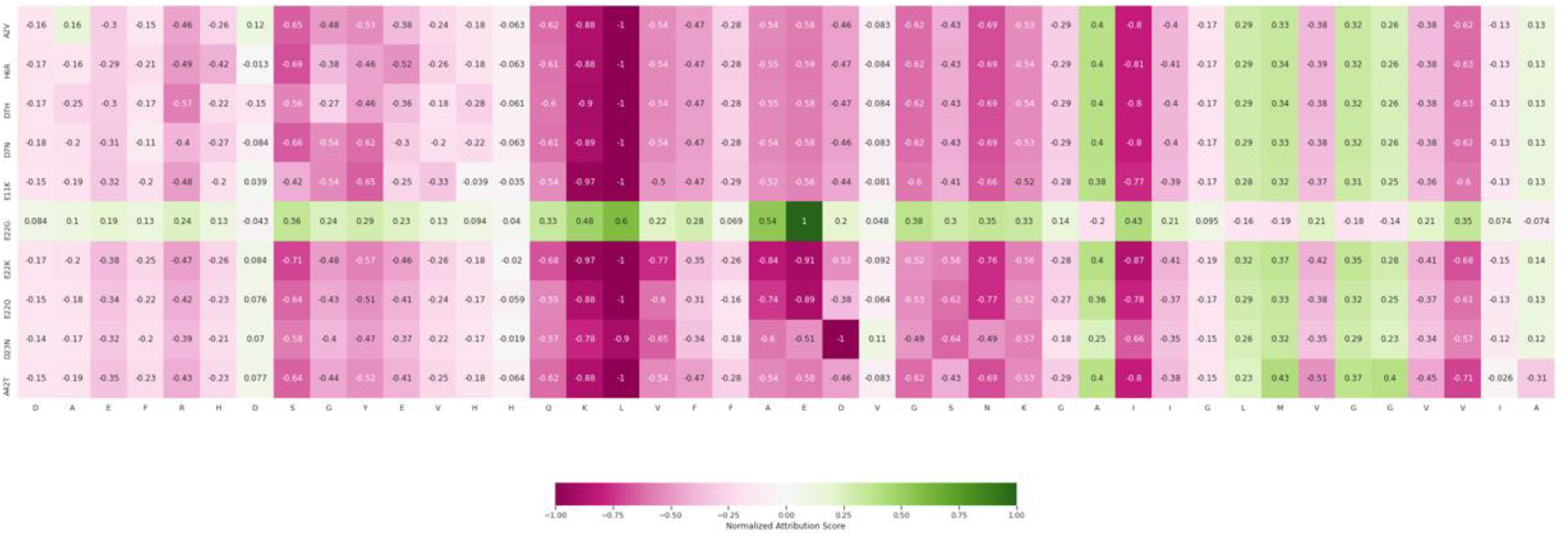
Normalized feature attribution from CNN-1D model. The Aβ_16-22_(KLVFFAE) fragment that is critical for self-assembly is highly relevant to the neural network, based on feature attribution from “Integrated Gradients”, in making a prediction. Important residues identified through Graph Operations (Figure 3C), including E3, V36, V39, were also notable in their feature attribution in the sequence-based CNN-1D model, indicating a close relationship between the attribution of prediction performance to input features and the Graph Operations based approach to inferring key features from empirically-observed protein structures.

## Discussion

In this work, we implemented seven neural networks to investigate the sequence-structure-phenotype relationship for Aβ_42_, a well-known pathogenic peptide for AD. The study of mutational effects is a long-standing research interest in a wide range of disciplines from biophysics, biochemistry, molecular biology, and genetics. Machine Learning / Deep Learning methods continue to be active areas of research in studies of protein engineering^18^ and annotations^22^ due to their high prediction accuracy and computational efficiency. For instance, with the aid of external descriptors, ConvNets models are able to extrapolate and explore novel mutations, thus overcoming the limitations of limited sampling from experiments. In our study, we constructed a comprehensive NN-derived mapping trained on DMS measurements, addressing the limited sampling (missing values) and the experimental noise (technical confounders). We have previously reported that external features such as an Amino Acid Based Descriptor (representing the intrinsic physicochemical properties of the amino acids) can enhance the prediction. Here, we showed that network architectures play a critical role. For instance, LR and FCN performed well only for the phenotype with the largest training data whereas ConvNets and RNN-LSTM showed superior performance across all three phenotypes even with a simple Identity Descriptor.

Using the prediction performance, we also exploited the neural networks as an inference tool to gain biological insights into mutational effects. As we showed, the network architecture implementation and the choice of hyperparameters determine how information is processed through the neural network. In addition, a hypothesis on biological mechanisms can be tested by designing the network appropriately or by leveraging the hyperparameters. In the case of sequence-based CNN, we found that applying dilated kernels can lead to degraded network performance. Although covering a larger convolutional window, kernels with high dilation rate neglect local information (motifs), which may be critical to the predicted phenotype (solubility). In our structure-based NN implementation, convolution directly operated on an irregular 3D grid of a GGA projected from the protein atomic coordinates. In our framework, which leverages the set of available, empirically-observed structures, model performance could be perturbed by both the input GGA and inter-residue distance threshold, allowing us to identify – via Graph Operations – residues and interactions that were critical for aggregation, i.e., to extract biological insights into relevant mechanisms. Similar approaches may open up new directions for utilizing neural networks towards other mechanistic studies, for example, analysis of protein stability.

Three aggregation-related phenotypes, curated from two independent DMS studies, were explored in this work. Nucleation is the bottleneck step along the aggregation trajectory whereas solubility is one of the important driving forces for inter-peptide interactions. While both studies aimed to leverage the phenotypes to elucidate the role of individual residues in Aβ aggregation, the aggregation pathways could differ^3^. Indeed, we found no strong correlation between the two NN-derived, mutationally-determined phenotypes (Figure S3), providing strong support for the presence of at least two aggregation processes. Notably, we found that the NN-derived nucleation provided significantly greater evidence than the original DMS phenotype for the critical roles of 7 gatekeeper residues, at which mutations would tend to increase nucleation. Finally, the NN-derived nucleation displayed improved performance relative to the original DMS phenotype in identifying known disease-causing mutations in Aβ_42_. Thus, an NN model of large-scale mutagenesis data may be leveraged as a powerful approach to fine-map the causal mutations in a disease-associated protein, with high computational efficiency and singular accuracy.

Our models built on protein sequences and DMS data may find application in genome-based systems. For example, Akita, a sequence-based CNN, has illustrated the power of neural networks in predicting 3D genome folding from sequence data^35^. As we showed here for protein data, GCN and RNN-LSTM models designed to reflect physicochemical and molecular insights may achieve improved prediction. Finally, we implemented IG to quantify the contribution of the input features to the neural network prediction. We found that inference from Graph Operations on empirically-observed protein structures was consistent with the feature attribution, providing confirmation of the utility of these operations in enabling explainability.

## Methods

### DMS datasets

We performed Deep Learning on publicly available DMS datasets^3,17^. Specifically, three physicochemical properties, i.e., solubility score (calculated from *Enrich2*)^17^, synonymous score^17^, and nucleation^3^ were curated.

### Engineering protein structural features

#### Generative Graph Architecture

Monomeric amyloid peptides are classified as intrinsically disordered proteins, although the later-stage aggregates present ordered conformations. We retrieved three structures from the Protein Data Bank with the following PDB IDs (1iyt^24^, 2nao^25^, and 5oqv^26^, Figure 3). Generative Graph Architectures (GGAs) were generated from the initial structural atomic coordinates, representing potential residue connections or interactions. Each residue was represented by a node while the presence of an edge was determined by whether the Euclidean distance (in ångströms) between a pair of residues (calculated from the coordinates of the corresponding *β*-carbons) was within a chosen threshold *d*_c_. Thus, chemically, the edges may encode intermolecular interactions, including covalent bonds, hydrogen bonds or van der Waals interactions between consecutive residues or non-consecutive residues. Varying the GGA input (from experimentally-obtained structures of the peptide) of a graph neural network – with the set of edges of the GGA for a given peptide structure also modified by varying the physical interaction radius *d*_c_ – allowed us to evaluate the contribution of specific residues (nodes) or interactions (edges) to the prediction performance.

#### Graph Operations

We operated on graph-objects/GGA via the NetworkX library^36^, from which we could easily acquire graph summary statistics, including number of nodes, number of edges, and centrality distribution. In our study, as all GGAs share the same number and ordering of the nodes (from the Aβ_42_ sequence), the GGA differences result from the edges. A GGA can also be mathematically defined in terms of its 2D adjacency matrix **M** where *M*_*ij*_ = 1 denotes a connection and *M*_*ij*_ = 0 denotes lack of connection for residues *v*_*i*_ and *v*_*j*_.

Graph operations are akin to *set operations*. For our purposes, they formalize operations on graphs to enable (downstream) evaluation of the contribution of specific nodes or edges to GCN model performance. The intersection *G*(*A, B*) of GGAs *A* and *B* returns the edges in both while the difference *H*(*A, B*) returns the edges present in GGA *A* only. Mathematically, the intersection *G*(*A, B*) defines an adjacency matrix **M** where the *ij*-th cell *M*_*ij*_ = 1 when *A*_*ij*_ = *B*_*ij*_ = 1 and 0 otherwise, with *A*_*ij*_ and *B*_*ij*_ denoting the *ij*-th cell of the adjacency matrices for the graphs *A* and *B*, respectively. Similarly, the difference *H*(*A, B*) defines an adjacency matrix **M** where *M*_*ij*_ = 1 when *A*_*ij*_ = 1, *B*_*ij*_ ≠ 1 and 0 otherwise. For example, the difference *H*(*A, B*) of GGAs *A* and *B* generated from the same peptide structure but using two different distance thresholds *d*_c,A_ and *d*_c,B_ (where, without loss of generality, *d*_c,A_ > *d*_c,B_) enables evaluation of the contribution of the edges unique to *A*. The adjacency matrices *A* and *B* as well as the derived matrices *G*(*A, B*) and *H*(*A, B*) are of equal dimension |*s*|^2^, where |*s*| is the length of the protein sequence *s* under consideration.

In addition to the adjacency matrix **M**, we define the degree matrix **D**, a diagonal matrix, with the *ii*-th cell given by the degree of the node *v*_*i*_, i.e., *D*_*ii*_ = deg (*v*_*i*_) (Figure 3C, for example), and the Laplacian matrix **L**:

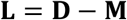

Note that *D*_*ii*_ = ∑_*j*_ *M*_*ij*_. The “influence” of a subset *A*_sub_ of nodes can be quantified by:

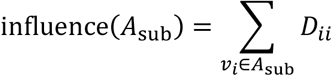

A connected component Ω is a subset of nodes such that any two nodes in Ω can be joined by a path of nodes entirely lying in Ω and there exist no paths between Ω and its complement Ω^*c*^. The number *φ* of connected components for the protein graph represented by **M** equals the vector space dimension of the kernel of the Laplacian **L**. If *λ*_1_, *λ*_2_, …, *λ*_*k*_ are the non-zero eigenvalues of **L**, then *φ* is given by:

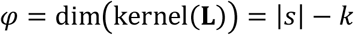

The connected components partition the GGA into disjoint subsets according to the interactions among the nodes (see Figure 3A, for example, where, at 6Å, the monomeric form forms a connected graph while both of the mature fibril conformations show multiple connected components).

### Network architectures and model training

In this work, we evaluated three classes of architectures, namely, Sequence-based: CNN-1D, CNN-2D, RNN-LSTM; Sequence and structure-based: GCN-AVE and GCN-SW; Baseline: FCN and LR. Four network architectures - CNN-2D, GCN-AVE, FCN and LR - were elements of a supervised learning framework previously implemented.^14,15^ CNN-1D, RNN-LSTM and GCN-SW were specifically tailored for the present study.

Network architectures utilized the Identity Descriptor (one-hot encoding) as well as the Amino Acid Based Descriptor (AAindex, encoding 566 intrinsic physicochemical properties of each amino acid) with the exception of LR and RNN-LSTM (which used the Identity Descriptor only). An RNN architecture, LSTM effectively comprehended local and global sequence information, learning context information. ProtCNN^22^ originated from ResNet^37^ and was designed for the problem of protein annotation/classification; in the implementation, after the initial convolution, the remaining two convolutional layers are packed within a residual block unit with a short-cut link. We adapted ProtCNN – constructing a model CNN-1D – so that it was feasible for the regression problem on DMS phenotypes. Similar to the CNN-2D model comprising of convolutional layers and one dropout layer as the last layer for regularization, a CNN-1D model was implemented to investigate the hyperparameters as well as interpret the information learned from the peptide sequence. CNN-1D is characterized by a simple architecture that utilized only one dilated convolutional layer with filter size set to 1. A dilated convolution is defined by:

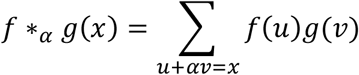

Here, α is the so-called dilation rate, with *α* ∈ ℕ and α≥1. When *α* = 1, the dilated convolution reduces to the conventional ConvNets.

GCN-AVE and GCN-SW were designed to integrate information from both sequence and structure inputs, directly operating on the GGAs instead of well-defined grids. In other words, edges in the GGAs were directly transformed into node connections in the neural network architecture. GCN-AVE was a variation of GCN-SW that distinguished center and neighboring nodes within the convolutional window, aiming to achieve better prediction for protein interface recognition^20^. On the other hand, GCN-SW had been developed for learning molecular fingerprints^21^, which made it an ideal model architecture to extract biological insights from structure-phenotype relationships.

Here we describe GCNs as applied to a GGA inherited from a peptide structure, given the importance of these architectures in the present study. Let **A** be the adjacency matrix of the GGA of a given peptide structure and *x*_*i*_ ∈ ℝ^ω^ be a feature description for the node *v*_*i*_. As before, let |*s*| be the peptide sequence length, which is also therefore the number of nodes. The input *X* to the GCN is a |*s*| x *ω* matrix, and the output *Y* is a node-level output of dimension |*s*| x *τ*. For instance, for the nucleation model, the output *Y* is a |*s*| x 1 matrix. The (*l* + 1)-st layer of a GCN (with *r* hidden layers) is a non-linear mapping:

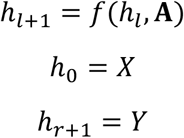

The non-linear mapping *f* can take on many forms, e.g., *h*_*l*+1_ = *σ*(**A***h*_*l*_*W*_*l*_), where *σ* is the activation function (e.g., leaky ReLU) and *W*_*l*_ is a matrix of weights. Instead of **A**, we can substitute **D**^−1^**A**, where **D** is the diagonal degree matrix (defined above), which would result in the averaging of the neighboring node feature description. The layers are permutation equivariant, resulting in the same *f* up to relabeling (permutation) of the nodes:

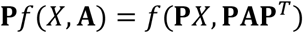

for a permutation matrix **P**. A specific GCN of interest is one defined by the following:

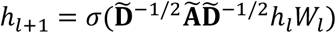

where **Ã** = **A** + **I** is the adjacency matrix of a graph with self-connections and 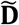 is the diagonal matrix defined by 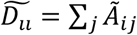.

We also implemented an RNN-LSTM. An RNN is particularly suited for sequence analysis. A hidden state at time step *t* + 1, denoted by *h*_*t*+1_, depends on both the current input and the previous hidden state:

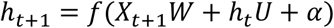

for an activation function *f* and bias *α*. The output at the time step *t* + 1, denoted by *O*_*t*+1_, is given by:

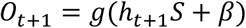

for an activation function *g* and bias *β*. The LSTM is a solution to the so-called vanishing gradient problem with RNN. An LSTM neuron in a recurrent layer controls the input and updates the neuron’s cell state and hidden state via ‘input’, ‘output’, and ‘forget’ gates.

Neural networks were trained using mean squared error as the loss function. We trained the models using the Adam^38^ optimizer (an extension of stochastic gradient descent), with dropout^39^ (rate=20%) as the regularization method to reduce overfitting. Leaky ReLU, which permits small negative values when the input values are negative, was applied as the activation function. Implementation was done using TensorFlow^40^. Additional implementation parameters (e.g., number of layers), which may vary with the model, are found in Table S1 and Table S3.

### Feature attribution

We utilized Integrated Gradients as a feature attribution method. For a given model *F*(*x*), the feature attribution for the *i*-th feature of the input vector *x* is defined as:

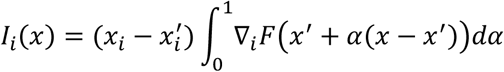

where *x*_*i*_ and 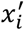 are the *i*-th component of *x* and the baseline *x*′, respectively, and ∇_*i*_*F* is the *i*-th component of the gradient of *F*. Here *α* can be interpreted as a perturbation quantity.

### Causal inference

Causal inference is a natural methodological consequence of deep learning applied to DMS data. As DMS experiments are frequently characterized by the presence of a large proportion of missing values and by the existence of possible confounding factors (affecting the phenotype), estimation of the mutationally-determined causal effects is an important problem to address. The “average mutational effect” *δ* on the phenotype *Y* is defined as follows:

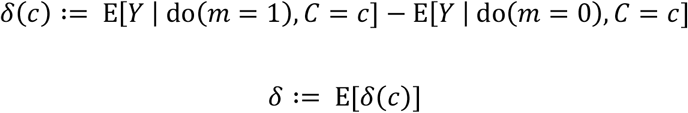

Here, *m* is an indicator for mutation at a residue (1=mutation, 0=wild-type), *C* is the set of additional covariates, and E is the expectation operator. The “do” operator follows Pearl’s notation. Our approach estimates *δ* using a neural network by estimating the conditional expectation E[*Y* | *m, C*] (Figure 4B) from a DMS-based sample {(*Y*_*i*_, *m*_*i*_, *C*_*i*_)}, drawn *i*.*i*.*d*. from the unknown probability distribution *P*, via the neural network prediction *Ŷ*. The average mutational effect *δ* can be used to prioritize residues or make inference on specific regions of the peptide (as done here on the difference in mutational effect between the C-terminus and N-terminus).

### Graph Operation-based perturbative approach to identify relevant features

The prediction performance for a given neural network model – quantified as the correlation cor(*Y, Ŷ*) between the phenotype *Y* and the predicted phenotype *Ŷ* – while the input GGA (with adjacency matrix **A**) is perturbed (e.g., via a Graph Operation or physical interaction radius thresholding on the atomic-resolution structure), is used to quantify the effect of the perturbation (such as, for illustration, the removal of specific edges to obtain a new adjacency matrix **A**_∗_):

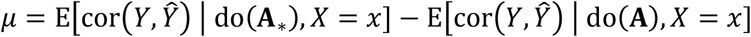

The statistic *µ* can be used on experimentally-obtained protein structures to evaluate the effect of inter-residue interactions and identify the specific interactions (and the implicated residues) that enhance the NN-derived, mutationally-determined phenotype. One advantage of this perturbative approach applied to Graph Operations is that it identifies relevant features – in a potentially complex combination rather than in isolation – by capturing their effects across empirically-observed structures.

### Performance

For each neural network model, the dataset was randomly split into 60% training, 20% tuning, and 20% test subsets. Trained models were evaluated for prediction performance on the test subset using Spearman correlation.

The phenotype *Y* and the NN-derived estimate *Ŷ* were shown above to be highly correlated, i.e., with *ρ* = cor(*Y, Ŷ*) = 1 − ε, for some small ε. Here, we calculate the probability of their difference in absolute value exceeding a given threshold *T, P*(|*Y* − *Ŷ*| > *T*). For any random variable *U* with mean *µ* and standard deviation *σ*, Chebyshev’s inequality states:

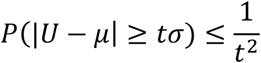

Hence, we obtain the following (after the substitution *U* = *Y* − *Ŷ*):

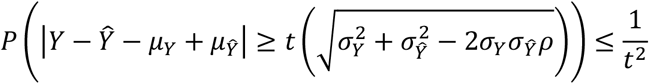

where *µ*_∗_ and *σ*_∗_ are the mean and the standard deviation, respectively, of the phenotype represented by the asterisk (observed or NN-derived). Assuming that *Y* and *Ŷ* are zero-centered with equal variance, the inequality reduces to:

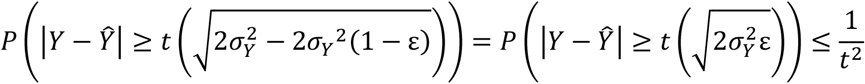

Equivalently, by setting 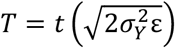, we obtain:

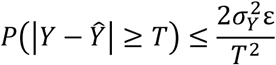

Thus, the worst-case scenario is determined here by the neural network performance (i.e., ε; the higher the performance, the smaller the ε) and the variability in the phenotype (i.e.,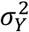).

## Data availability

All results are available in Supplementary Information.

## Code availability

The pipelines and configurations used in the analyses are available in the project’s Github repository: https://github.com/gamazonlab/DeepLearning_Abeta.

## Acknowledgments

E.R.G. is grateful to the President and Fellows of Clare Hall, University of Cambridge for providing a stimulating intellectual home. This research is supported by the following NIH grants: NHGRI R35HG010718, NHGRI R01HG011138, NIA AG068026 and NIGMS R01GM140287.

## Author contributions

E.R.G. and B.W. designed the study and wrote the manuscript. B.W. performed the experiments and analyses. E.R.G. conceived and directed the study.

