## Supplementary Information for "Towards mechanistic models of mutational effects: Deep Learning on Alzheimer’s Aβ peptide"

Bo Wang[1] and Eric R. Gamazon[1, 2, 3, 4]

[1]Division of Genetic Medicine, Department of Medicine, Vanderbilt University Medical Center, Nashville, TN, USA

[2]Vanderbit Genetics Institute, Vanderbilt University Medical Center, Nashville, TN, USA

[3]Data Science Institute, Vanderbilt University Medical Center, Nashville, TN

[4]Clare Hall, University of Cambridge, Cambridge, United Kingdom

Send correspondence to:

Eric R. Gamazon <>

**Table S1.** Neural network performance on $A\beta_{42}$ biochemical phenotypes. The phenotypes were "Nucleation", "Solubility", and "Synonymous". Performance for each phenotype was evaluated using the Spearman correlation between predicted and observed phenotype in a test set (independent of the training set). PDB refers to the Protein Data Bank identifier for the protein structure.

| Neural networks | Nucleation | Solubility | Synonymous | PDB | Model Architecture |
|---|---|---|---|---|---|
| CNN-1D | 0.88557 | 0.84072 | 0.82354 | - | Layer: 3 Kernel:5 Filter:32 |
| CNN-2D | 0.87945 | 0.90599 | 0.89683 | - | Layer: 3 Kernel:5 Filter:32 |
| RNN | 0.87496 | 0.90969 | 0.90608 | - | Bidirectional LSTM |
| GCN-AVE, threshold 9 | 0.88036 | 0.92124 | 0.92327 | 1iyt | Layer:2 Filter:32 |
| GCN-SW, threshold 6 | 0.73519 | - | - | 1iyt | Layer:1 Filter:1 |
| GCN-SW, threshold 14 | 0.73235 | - | - | 1iyt | Layer:1 Filter:1 |
| GCN-AVE, threshold 9 | 0.88140 | 0.91905 | 0.93373 | 2nao | Layer:2 Filter:32 |
| GCN-SW, threshold 6 | 0.59761 | - | - | 2nao | Layer:1 Filter:1 |
| GCN-SW, threshold 14 | 0.74120 | - | - | 2nao | Layer:1 Filter:1 |
| GCN-AVE, threshold 9 | 0.882756 | 0.92694 | 0.92929 | 5oqv | Layer:2 Filter:32 |
| GCN-SW, threshold 6 | 0.63552 | - | - | 5oqv | Layer:1 Filter:1 |
| GCN-SW, threshold 14 | 0.64655 | - | - | 5oqv | Layer:1 Filter:1 |
| LR | 0.86213 | 0.57034 | 0.38316 | - | - |
| FCN | 0.85837 | 0.71882 | 0.57034 | - | Layer:1 Nodes:100 |

**Table S2.** Performance of one-layer CNN-1D at various choices of kernel size and dilation rate. Performance for each phenotype was evaluated using the Spearman correlation between predicted and observed phenotype in a test set (independent of the training set).

| Dilation Rate | Kernel | Modified Kernel | Performance | Stdev |
|---|---|---|---|---|
| 1 | 5 | 5 | 0.79835 | 0.02912 |
| 2 | 3 | 5 | 0.75789 | 0.03722 |
| 1 | 6 | 6 | 0.86151 | 0.02081 |
| 1 | 7 | 7 | 0.82246 | 0.02396 |
| 2 | 4 | 7 | 0.83643 | 0.0252 |
| 3 | 3 | 7 | 0.75379 | 0.03312 |
| 1 | 9 | 9 | 0.83892 | 0.02234 |
| 2 | 5 | 9 | 0.72694 | 0.0371 |
| 4 | 3 | 9 | 0.42581 | 0.04859 |

**Table S3.** Additional hyperparameters for the neural network models

| Models | Patience | Maximum Number of Epochs | Training |
|---|---|---|---|
| GCN-AVE, GCN-SW, CNN-2D | 10 | 400 | Learning rate: 0.0001, Batch size: 128, Dropout rate: 0.2, Activation: 'leaky_relu' |
| CNN-1D, RNN-LSTM | 30 | 2000 | |

*For single layer CNN-1D, residue block was removed, padding = ‘causal’, activation=’None’

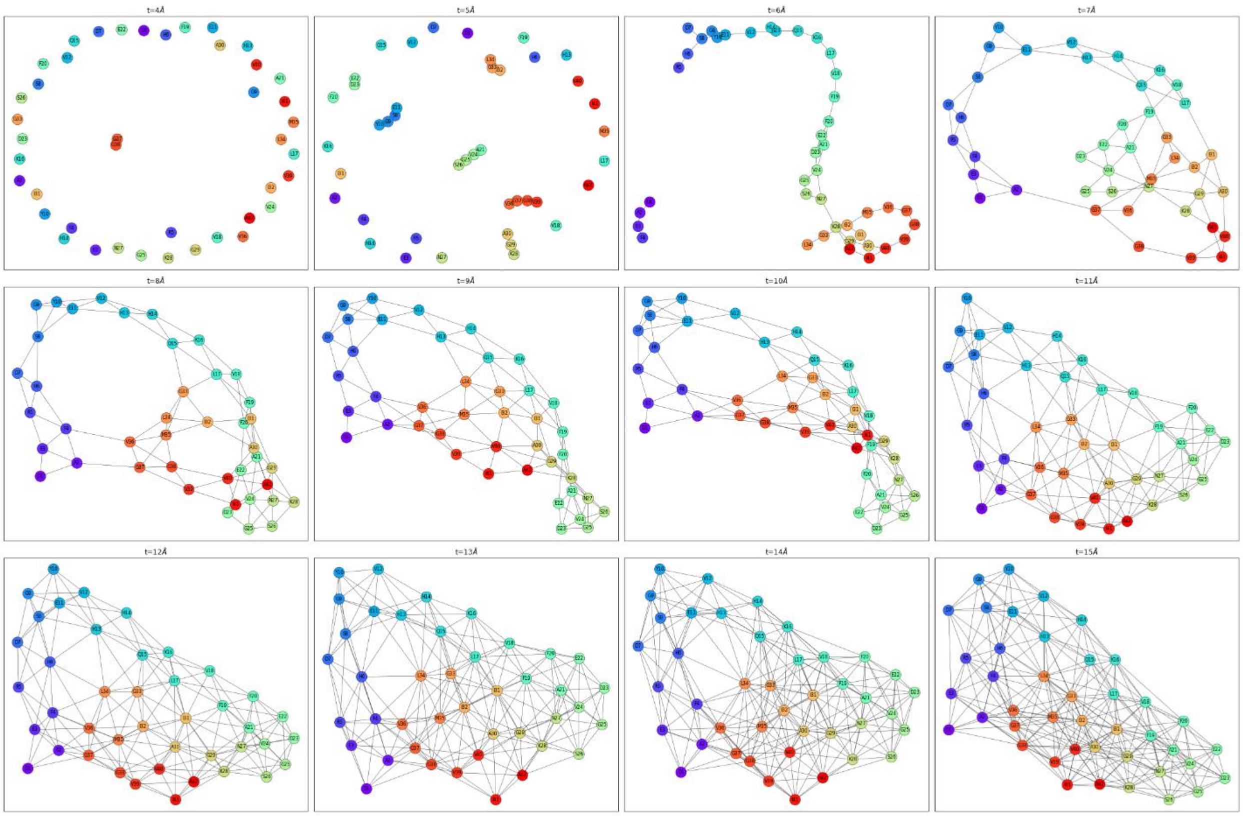


**Figure S1. Example GGAs for structure with PDB ID 2nao from varying the distance threshold with values ranging from 4Å to 15Å.**

A

B

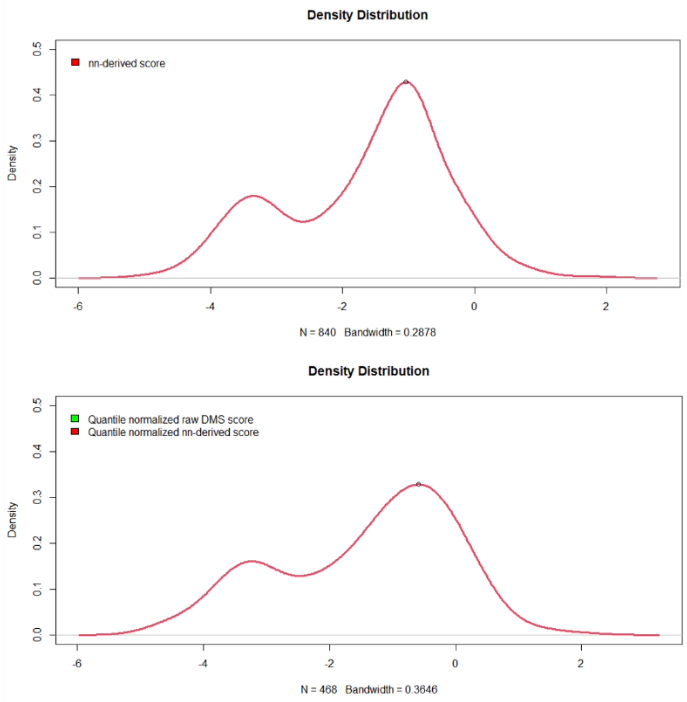


**Figure S2. Quantile normalization on Raw DMS measurement and NN-derived score.** The position of the right peak indicates the adjusted threshold (from 0 to -0.5844597) used to discriminate the nucleation effect. In **B**, the two curves are directly overlapping.

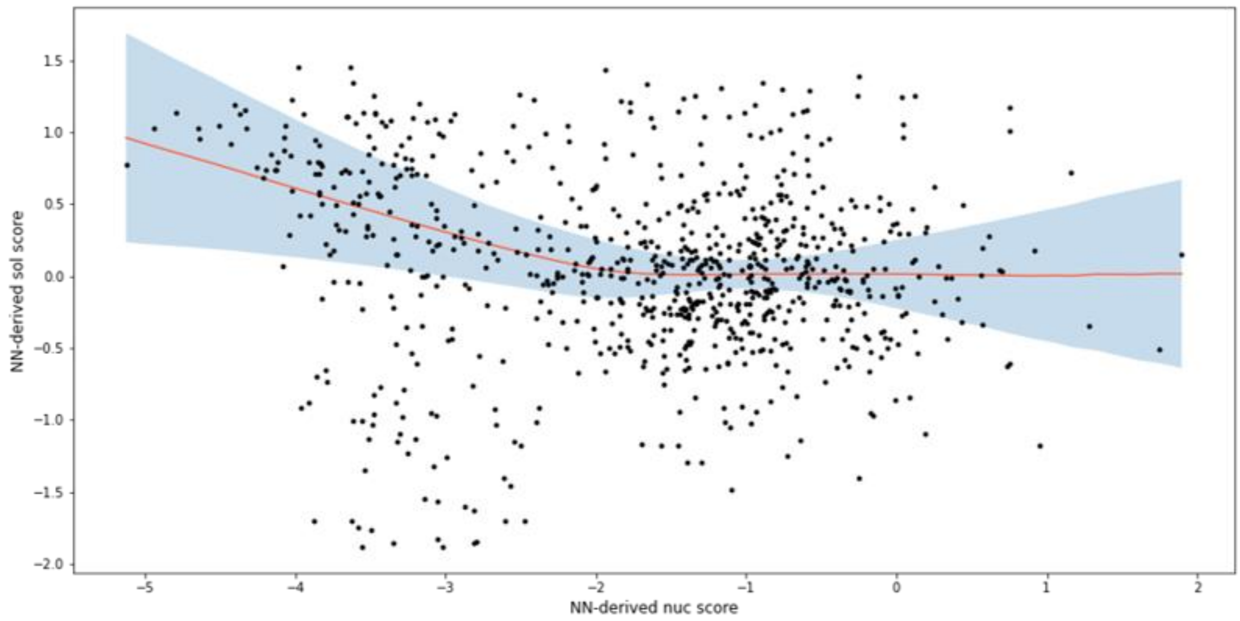


**Figure S3. Relationship between two NN-derived phenotypes: nucleation score and solubility score.** We fit a curve through the points, using Locally Estimated Scatterplot Smoothing (LOESS), a method for local polynomial regression.
